# A Combinatorial Strategy for HRV 3C Protease Engineering to Achieve the N-terminal Free Cleavage

**DOI:** 10.1101/2024.01.05.574269

**Authors:** Meng Mei, Xian Fan, Yu Zhou, Faying Zhang, Guimin Zhang, Li Yi

## Abstract

Human rhinovirus 3C protease (HRV 3C-P) has a high specificity against the substrate of LEVLFQ↓G at P1’ site, which plays an important role in biotechnology and academia as a fusion tag removal tool. However, a non-ignorable limitation is that an extra residue of Gly would remain at the N terminus of the recombinant target protein after cleavage with HRV 3C-P, thus potentially causing protein mis-functionality or immunogenicity. Here, we developed a combinatorial strategy by integrating structure-guided library design and high-throughput screening of eYESS approach for HRV 3C-P engineering to expand its P1’ specificity. Finally, a C3 variant was obtained, exhibiting a broad substrate P1’ specificity to recognize 20 different amino acids with the highest activity against LEVLFQ↓M (*k_cat_*/*K_M_*= 3.72 ± 0.04 mM^-1^·s^-1^). Further biochemical and NGS-mediated substrate profiling analysis showed that C3 variant still kept its substrate stringency at P1 site and good residue tolerance at P2’ site, but with an expanded P1’ specificity. Structural simulation of C3 indicated a reconstructed S1’ binding pocket as well as new interactions with the substrates. Overall, our studies here prompt not only the practical applications and understanding of substrate recognition mechanisms of HRV 3C-P, also provide new tools for other enzyme engineering.

**Abstract graphic:** 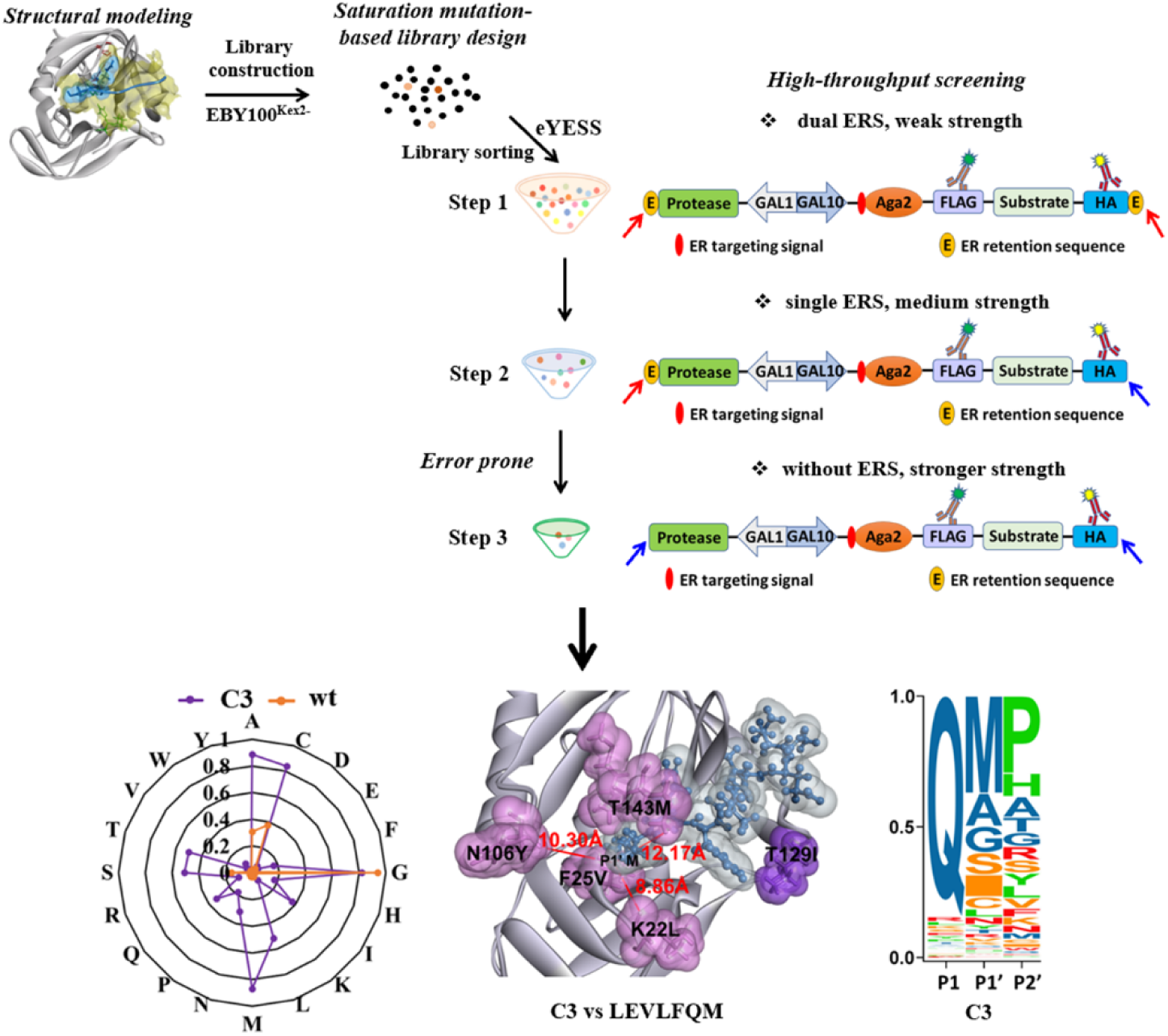

## 1. Introduction

Proteases play important roles in biology, including the initiation, regulation, and termination of physiological processes[1–3]. The unique ability of protease to catalyze reactions with exquisite specificity makes them useful in therapeutic and biotechnological applications. The US Food and Drug Administration has approved more than 12 therapeutic proteases, including anticoagulants tPA and uPA, Factor IX, and Thrombin, etc[4]. In biotechnological applications, proteases are widely used for investigation of protein-protein interactions[5, 6], synthetic sensor-and-response circuits in signal transduction[7–9], and affinity purification tag removal[10–12].

Many proteases have low substrate specificities, such as trypsin, Factor Xa, papain, etc. In comparison, viral proteases, such as Tobacco Etch Virus (TEV) protease and human rhinovirus (HRV) 3C protease have high substrate specificities. In general, strict substrate specificity of protease confers advantages of specific cleavage, thus favoring their practical applications through producing uniform products or providing precise regulation. For example, TEV protease and HRV 3C protease have been commonly used in fusion tag removal in the protein production. However, their high substrate specificity causes extra residues left to the target proteins after proteolytic cleavage, which might cause function lose or immunogenicity of the target proteins. Therefore, engineering proteases against new substrates but still with high proteolytic activity would greatly expand their applications in therapeutics and academics. Although it is a very challenging task to engineer highly specific protease, several impressive progresses have been recently reported using directed evolution incorporated with newly developed high-throughput screening methods[13–17]. For example, yeast endoplasmic reticulum (ER) sequestration screening (YESS) system was developed to successfully engineer the TEV protease to recognize substrates with different P1 residues[18]. Sanchez and Ting engineered the TEV protease to improve its activity in a yeast platform using the concept of proteolytic release of a membrane-anchored transcription factor[15]. Packer and Liu engineered TEV protease with PACE system to efficiently cleave a very different sequence of HPLVGHM in human IL-23[13]. Carrico and Francis combined selection and counter-selection strategy via antibiotic resistance to obtain a TEV protease mutant with new specificity at P6 residue[19]. Similar to TEV protease, HRV 3C protease (HRV 3C-P) is also a type C protease, belonging to the cysteine protease family with the catalytic triad of His_40_-Glu_71_-Cys_146_. HRV 3C protease recognizes the peptide sequence LEVLFQ↓G with very high specificity at P1 and P1’ residues[20]. Its most extraordinary characteristics is that HRV 3C protease presents high activity in a broad temperature range from 4 to 30 [[21–23]. Because of its robust activity at low temperature of 4 [, HRV 3C protease is widely used as an effective fusion tag removal tool, especially for those cryogenic proteins or therapeutic proteins whose purification need to be carried out at low temperature. The major limitation of its practical application is that HRV 3C-P has a high substrate specificity at the P1’ site of Gly[20], which leads an extra Gly at the N-terminus of recombinant target protein to potentially affect its practical application. However, compared to TEV protease, no study has been reported on the directed evolution of HRV 3C protease. Since Met is the starting residue for most proteins, engineering HRV 3C-P against LEVLFQ↓M could confer non-modification of the N-terminus of recombinant target protein. Even more, expanding its substrate specificity to recognize all 20 amino acids at the P1’ site could expand its applications with the ability of completely cleaving the N-terminal fusion tags of the target protein regardless of the first amino acid.

Yeast surface display (YSD) system combined with the fluorescence activated cell sorting (FACS) technology has been proven as a powerful high-throughput screening approach used in protein directed evolution[24]. Our previously established yeast endoplasmic reticulum (ER) sequestration screening (YESS) approach was one of the pioneer high-throughput protease engineering platforms, which combined YSD, FACS technology, yeast ER protein retention, and directed evolution to effectively carry out single-cell high-throughput screening of TEV protease libraries[18]. In YESS system, the C-terminal ER retention sequence (ERS) is used to retain protease and its substrates in the yeast ER to regulate the method sensitivity. In principle, the strong ERS would prolong their residence within the yeast ER, thus prompting the slow protease hydrolysis reaction to amplify the fluorescence signal for detection. In contrary, the weak ERS and no ERS will decrease the residence of protease and its substrates within the yeast ER to help differentiate the subtle difference of fast protease hydrolysis reactions. Through this approach, YESS has engineered TEV protease for altered substrate P1 specificity, obtaining highly active variants with 5,000-fold and 1,000-fold specificity alteration for Glu or His at the substrate P1 site, respectively[18].

Here, an enhanced YESS (eYESS) approach was developed to provide adjustable sensitivity to quantify weak protease hydrolysis signals and differentiate fast proteases with subtle catalytical ability difference. The LEVLFQ↓M substrate is used as the substrate for the HRV 3C-P engineering to expand its substrate p1’ specificity. Finally, an engineered C3 variant was obtained with a *k_cat_*/*K_M_* value of 3.72 ± 0.04 mM^-1^·s^-1^ against LEVLFQ↓M. The further characterization of C3 variant indicated that it had a re-constructed S1’ pocket to recognize all 20 residues at the P1’ site.

## 2. Materials and methods

### 2.1. Vector construction

Based on previously published pESD vector[25], the HRV 3C-P (GI: 25121860) and its substrates LEVLFQ↓X (X can be any residues) encoding gene were cloned to *Pst*I/*Eco*RI sites and *Bam*HI/*Xho*I sites, respectively, to generate a HRV 3C-P-*GAL1*-*GAL10*-Aga2-FLAG-LEVLFQ↓X-HA complex cassette. Different ERSs were anchored at the C-terminus of both HRV 3C-P and Aga2-FLAG-LEVLFQ↓X-HA cassette. The wt HRV 3C-P or its variants gene was cloned into the pRK792 vector to generate MBP-LEVLFQ↓X-6His-HRV 3C-P complex cassette for protein expression. Meanwhile, different protein substrates including MBP-LEVLFQ↓X-6His-GST and MBP-LEVLFQ↓X-6His-GFP cassettes were also cloned into the pRK792 vector for *in vitro* proteolytic activities experiments of wt HRV 3C-P and its variants. Detailed information is listed in supplemental Table S1.

### 2.2. Protease library construction

Fragments including HRV 3C-P gene, ERS (FEHDEL) encoding region, and 40 bp homologous recombination sequence were assembled by overlapping PCR to generate a saturation mutagenesis DNA library of six residues including Lys22, Phe25, Asn106, Thr143, Gly144, and Gln145 in the S1’ pocket of HRV 3C-P. The library was generated using saturation mutation with the combination of NNS and NDT, VMA, ATG, TGG degenerated codons (N is any nucleotide, S = G/C, D = A/G/T, V = A/C/G, and M = A/C). In addition, the four degenerated codons of NDT/VMA/ATG/TGG covering 20 amino acids could effectively reduce the codon redundancy of the library. For the first step of screening, the HRV 3C-P DNA library was inserted downstream of the *GAL1* promoter in the pYSD-1 vector (**Table S1**), containing the Aga2-FLAG-LEVLFQ↓M-HA-FEHDEL complex cassette downstream of the *GAL10* promoter. Prepared HRV 3C-P library DNA and linearized pYSD-2 vector were mixed at a molar ratio of 3:1 and then transformed into *Saccharomyces cerevisiae* strain EBY100^Kex2-^ electrocompetent cells[26]. Library transformation and cell cultivation were performed with reference to the previous published method[27].

To remove the ERS (FEHDEL) at the C-terminus of HRV 3C-P library DNA in the second step of screening, plasmids were extracted from sorted cells and HRV 3C-P library DNA was amplified. Amplified HRV 3C-P library DNA was mixed with linearized pYSD-2 vector (**Table S1**) containing the Aga2-FLAG-LEVLFQ↓M-HA complex cassette downstream of the *GAL10* promoter, and then transformed into *S. cerevisiae strain* EBY100^Kex2-^ electrocompetent cells as above method, generating the second library for further sorting. Finally, a random mutagenesis library based on HRV 3C-P variant (B22) without ERS (FEHDEL) was generated by error-PCR amplification (1.0-1.5% error rate), and then inserted downstream of the *GAL1* promoter in the pYSD-2 vector (**Table S1**) in the third step of screening.

### 2.3. FACS analysis and yeast cell screening

Transformed yeast cells containing HRV 3C-P or its variants gene and their substrate complex cassettes were cultivated in YNB-Glucose-CAA medium to an OD_600_ of 2.0-3.0, followed by induction with an initial OD_600_ of 0.5 in YNB-Galactose-CAA medium. After induction for 8 h at 30[, yeast cells were harvested by centrifugation, washed and incubated with 0.1 μM anti-HA-FITC antibody and 0.1 μM anti-FLAG-iFluor 647 antibody (GenScript, Nanjing, China) for 15 min in dark. Subsequently, cell fluorescence analysis was detected with the FITC channel (525/40 nm band pass) and APC channel (660/20 nm band pass) by using Beckman Coulter CytoFLEX (Brea, CA) equipped with 488 and 633 nm lasers, respectively. The proteolytic efficiencies of HRV 3C-P and its variants were calculated as: [Normalized proteolytic efficiency] = [The percentage of cells with only iFluor 647 fluorescence signals] / ([The percentage of cells with only iFluor 647 fluorescence signals] + [The percentage of cells with both iFluor 647 and FITC fluorescence signals]).

∼2 × 10^9^ cells bearing HRV 3C-P libraries, around 10-fold larger than the designed library size, were grown, induced and labeled as above method. The antibody-labeled cells were resuspended in 1× PBS buffer and sorted using Beckman Coulter MoFlo XDP flow cytometer (Brea, CA, USA) equipped with the FITC channel (525/40 nm band pass) and APC channel (660/20 nm band pass). In each round, 10^6^-10^8^ cells with the high iFluor 647 and low signals were sorted. Collected cells were grown in YNB-Glucose-CAA medium and induced in YNB-Galactose-CAA medium for the next round of sorting. After 3 rounds of sorting, aliquots of collected cells were spread on YNB-Glucose-CAA plates and 50 single clones were reanalyzed by FACS analysis with the above method. Yeast plasmids containing HRV 3C-P variant gene were extracted by using Zymolyase (Amsbio, Abingdon, UK) and transformed into *E. coli* XL-Gold for plasmid rescue, and then were sequenced to obtain the mutation information of HRV 3C-P variants.

### 2.4. Protein expression and purification of HRV 3C-P or its variants and their substrates

Plasmids bearing HRV 3C-P or its substrate complex cassettes were transformed into *E. coli* BL21 (DE3) cells, respectively. The cells were grown to an OD_600_ of 0.6-0.8 in LB medium at 37 [, and then induced with 0.5 mM IPTG at 18 [. After induction for 20 hours, cells were collected and resuspended in lysis buffer containing 50 mM Tris-HCl (pH 8.0), 200 mM NaCl, 50 mM NaH_2_PO_4_, 0.5 mg/mL lysozyme, 1 mM PMSF, 10% glycerol, 25 mM imidazole, and 5 mM β-mercaptoethanol for sonication. The supernatant containing HRV 3C-P or its substrates was obtained by centrifugation at 10,000 × g at 4 [for 20 min, and was then subjected to Ni-NTA affinity purification at 4 [(Qiagen, Valencia, CA). The column was adequately washed with 20 column volume of wash buffer containing 50 mM Tris-HCl (pH 8.0), 200 mM NaCl, 50 mM NaH_2_PO_4_, 40 mM imidazole, and 5 mM β-mercaptoethanol. The protein was eluted with elution buffer containing 50 mM Tris-HCl (pH 8.0), 200 mM NaCl, 50 mM NaH_2_PO_4_, 200 mM imidazole, and 5 mM β-mercaptoethanol, and was then desalted with desalting column (GE Healthcare, Chicago, IL, USA) using AKTA system (GE Healthcare, Chicago, IL, USA). Finally, the purity and concentration of purified protein were analyzed by SDS-PAGE and UV absorbance measurement, respectively.

### 2.5. Protein substrate cleavage analysis of HRV 3C-P and its variants

Purified wt HRV 3C-P or its variants were mixed with the purified substrate MBP-LEVLFQ↓X-6His-GST at different molar ratios based on different experimental requirements. The proteolytic activities of HRV 3C-P or its variants were measured in total 200 μL cleavage buffer containing 50 mM Tris-HCl (pH 8.0), 150 mM NaCl, and 5 mM mercaptoethanol. After reaction at 4 [for 1 h or 4 h, proteolysis was terminated by mixing the reaction sample with SDS–PAGE protein loading buffer and boiled at 100 °C for 10 min, followed by SDS-PAGE analysis. The proteolytic efficiency was quantitated based on the change of band intensities and calculated as: [Proteolytic efficiency] = (1-[Band intensity of uncut protein substrate MBP-LEVLFQ↓X-6His-GST] / [Band intensity of the control of protein substrate MBP-LEVLFQ↓X-6His-GST]) × 100%.

### 2.6. HRV 3C-P kinetic assays

Different concentration of purified protein substrates MBP-LEVLFQ↓X-6His-GFP were incubated with 100 nM or 500 nM purified protease (wt HRV 3C-P and C3 variant) based on different experimental requirements. 100 nM purified protease (wt HRV 3C-P and C3) were incubated with substrate MBP-LEVLFQ↓G-6His-GFP or MBP-LEVLFQ↓M-6His-GFP at different concentrations of 5 µM, 15 µM, 30 µM, 60 µM, 120 µM, and 320 µM. Each reaction was performed in total 200 μL cleavage buffer at 4 °C. For each reaction, the samples were taken at 6 different time points 0, 5, 10, 15, 25 and 45 min, and were terminated by mixing the reaction sample with SDS–PAGE protein loading buffer, followed by immediately flash freezing in liquid nitrogen. The proteolysis was analyzed by SDS–PAGE at 4 °C and in-gel fluorescence using Amersham™ Typhoon instrument (Cytiva, Marlborough, MA). Band intensities were measured and fit to a Michaelis–Menten enzyme-kinetics model to calculate the kinetic parameters. Due to the low solubility of MBP-LEVLFQ↓X-6His-GFP, we calculated the initial catalytic rate at the maximum concentration of 360 μM for the substrates with low cleavage efficiency.

### 2.7. Structure modeling

The peptide substrates were obtained based on the published peptide structure covalently bound to the wt HRV 3C-P (PDB: 2B0F[28]). The structures of HRV 3C-P variants were simulated by AphaFold 2[29], followed by docking with different peptide substrates through the ZDOCK program in Discovery Studio 2021 with reference to the previous published method[20].

## 3. Results and discussion

### 3.1. A combinatorial strategy for HRV 3C protease engineering

Currently, protein engineering has evolved from the early random or site-mutagenesis to combinatorial methods, which normally incorporate structure-guided and computational library design, and directed evolution with high-throughput screening[24, 30, 31]. The first generation YESS system has proved to be a facile and general strategy for engineering protease with altered substrate specificity and catalytic activity[18]. Here, we established an enhanced YESS approach (eYESS), in which 3 steps were conducted for the directed evolution of HRV 3C protease to alter its substrate P1’ specificity (**Figure 1A**). FEHDEL was used both at the C terminus of protease library and its substrate in the first step to enrich enough protease variants with either high or low activity. Once these variants were enriched, in the second step, the ERS at the C-terminus of substrate was removed to increase the screening stringency, thus enriching protease variants with moderate improved activity. In the third step, the ERS at the C-terminal of protease was also removed to further increase the screening stringency, and only fast protease variants with the high proteolytic activity could be isolated. Moreover, to increase the effectiveness of designed library, a Kex2 knockout yeast strain EBY100^Kex2-^ was used for library screening, which could largely prevent the target protease and its substrate from being cleaved by the major endogenous protease Kex2 in the yeast secretory pathway[26].

ERS is a key element that plays a critical role in the eYESS approach. By varying the ERS, different sorting stringency can be generated to provide a more precisely controlled manner for library screening. The function of the ERS was also confirmed in another recently established yeast-based high-throughput protein engineering approach, the PrECISE (protease evolution via cleavage of an intracellular substrate) system[32]. Also, a tunable platform by incorporating modulate gene transcription, YESS 2.0 system, was recently designed to further improve a TEV-P variant with a 2.25-fold higher catalytic efficiency[16].

**Figure 1.**
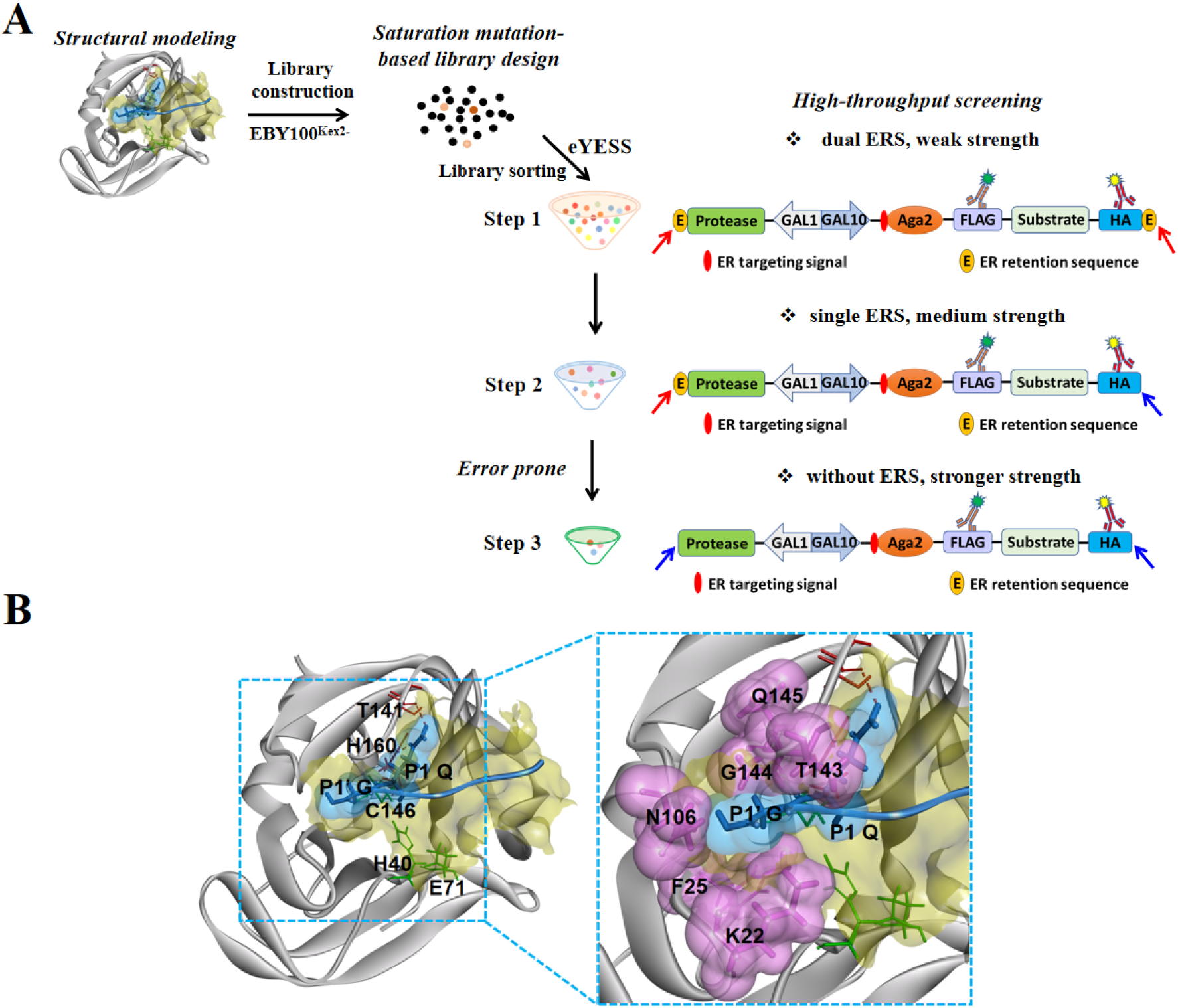
The combinatorial strategy for HRV 3C protease engineering. A. Overview of the 3 steps of eYESS strategy for protease engineering. B. Molecular docking of HRV 3C-P with LEVLFQ↓G. The yellow area shows the substrate binding pocket of HRV 3C-P (gray). The substrate LEVLFQ↓G is shown in blue, and Gln at P1 site and Gly at P1’ site are shown in blue spheres. The catalytic triplets His40-Glu71-Cys146 are indicated in green. In the S1 binding pocket, two H-bonds (red) are formed between Thr141 and His160 with Gln at the substrate P1 position (left). In the S1’ binding pocket, 6 residues consisting of Lys22, Phe25, Asn106, Thr143, Gly144, and Gln145 within 3.5 Å distance from Gly at substrate P1’ position are labeled in pink spheres (right).

### 3.2. Directed evolution of HRV 3C protease for altered P1’ specificity

In our previous studies, HRV 3C protease exhibited very high specificity at substrate P1 and P1’ sites, only recognizing Gln or Glu at the P1 position, and Gly, Ala, Ser, or Cys at the P1’ position even under the condition of using strong ERS of FEHDEL to prompt ER retention effect[20]. For the initial HRV 3C-P engineering for altered P1’ specificity, structural modeling and docking of HRV 3C-P with LEVLFQ↓G was performed through the ZDOCK program in Discovery Studio, based on the published HRV 3C-P structure (PDB: 2B0F[28]). As shown in **Figure 1B**, Thr141 and His160 in the S1 binding pocket of HRV 3C-P interacted with the substrate P1 position Gln through strong H-bonds to maintain its correct orientation. This orientation facilitated catalytic residue Cys146 to attack the Q_P1_-G_P1’_ peptide bond. Six residues consisting of Lys22, Phe25, Asn106, Thr143, Gly144, and Gln145 locate within 3.5 Å distance from Gly at substrate P1’ position, forming a very rigid and narrow S1’ binding pocket of HRV 3C-P to only recognize and hold the small residues at the P1’ position. Therefore, to re-construct a new S1’ binding pocket, a mutant library with a size of 2.7×10^8^ covering six residues (Lys22, Phe25, Asn106, Thr143, Gly144, and Gln145) in the S1’ binding pocket was generated to screen the variants against the substrate LEVLFQ↓M.

In the first step, FEHDEL was anchored at the C-terminus of both HRV 3C-P and Aga2-FLAG-LEVLFQ↓M-HA substrate cassette to provide a weak sorting stringency. The HRV 3C-P mutant library covering 6 mutation sites with the C-terminal ERS of FEHDEL was amplified, followed by yeast transformation together with the linearized pYSD-1 plasmid (**Table S1**) to obtain the naïve library. Approximately 2 × 10^9^ yeast cells were initially sorted through 3 consecutive rounds of FACS against high iFluor 647 (anti-FLAG-iFluor 647) and low FITC (anti-HA-FITC) signals, and ∼10^7^ yeast cells (named sub-library 1) were collected (**Figure S1**). Subsequently, 50 single clones from the sorted cells were picked up and characterized to evaluate the quality of the sorted sub-library 1. 20 variants show proteolytic activity against LEVLFQ↓M (**Table S2**), among which the A3 variant containing the K22M, N106T, T143P, and Q145T mutations showed the highest cleavage efficiency of 84.2% with double ERSs (**Figure S1**).

Next, HRV-3C-P protease genes with ERS of FEHDEL were amplified from the sorted sub-library 1 and re-transformed into yeast cells along with the linearized vector pYSD-2 (**Table S1**), in which FEHDEL is removed from C-terminus of Aga2-FLAG-LEVLFQ↓M-HA substrate cassette to increase the sorting stringency. In this step, 3 rounds of cell sorting were performed to enrich cells (names sub-library 2) displaying the high iFluor 647 but low FITC fluorescence. Again, 50 single clones from sub-library 2 were picked up and further analyzed. 10 variants showed proteolytic activity against LEVLFQ↓M (**Table S2**), among which the B22 variant containing the K22L, F25V, N106Y, T143M, and Q145M mutations showed the highest cleavage efficiency of 98.4% with single ERS (**Figure S1**).

However, it was noticed that the cleavage efficiency of B22 variant against LEVLFQ↓M decreased from 98.4% to 82.8% when the ERS sequence from the B22 variant construct was removed (**Figure S1**). Hereafter, in the third step, a random mutagenesis library based on the B22 variant sequence was generated by error-prone PCR (1.0-1.5% error rate), which was integrated into the vector pYSD-2 (**Table S1**). Finally, 3 rounds of library sorting were performed against ∼1 × 10^8^ yeast cells, and ∼10^6^ yeast cells displaying the high iFluor 647 but very low FITC fluorescence were collected (names sub-library 3). 50 single clones from sub-library 3 were further analyzed, among which two variants, C3 and C9, containing an additional T129I and A140S mutation based on the B22 (**Table S2**), respectively, were identified. Finally, our results indicated that C3 has the highest proteolytic activity against substrate LEVLFQ↓M with a cleavage efficiency of 98.6% without ERS (**Figure S1**).

### 3.3. The activity of HRV 3C-P variants against the protein substrate in vitro

To understand the catalytic mechanism of HRV 3C protease, A3, B22, and C3 variants from the sub-library 1, 2, and 3, respectively, along with wild-type (wt) HRV 3C protease and their protein substrates were recombinantly expressed using pRK-1 (MBP-LEVLFQ↓G/M-6His-GST) and pRK-3 (MBP-LEVLFQ↓G/M-6His-protease) vectors (**Table S1**) in *E.coli* cells and their proteolytical activity against different protein substrates was evaluated, respectively (**Figure 2A**). In principle, once the 70 kDa fusion protein substrate (MBP-LEVLFQ↓G/M-6His-GST) was cleaved by HRV 3C-P or its variants, it could be splitted into two parts (MBP-LEVLFQ (43 kDa) and G/M-6His-GST (26 kDa)), which can be analyzed through SDS-PAGE. As shown in **Figure 2A**, wt HRV 3C protease presented the highest cleavage efficiency of 72.9% against the MBP-LEVLFQ↓G-6His-GST substrate in a protease/substrate molar ratio of 1:50 at 4 [for 1 hour, but it showed no cleavage against the MBP-LEVLFQ↓M-6His-GST substrate. Comparably, under the same condition, A3, B22, and C3 variants all showed higher activities against the MBP-LEVLFQ↓M-6His-GST substrate than that of wt HRV 3C-P, with a cleavage efficiency of 4.9%, 56.2%, and 73.3%, respectively. Especially for the C3 variant, it also presented a high cleavage efficiency of 45.1% against the MBP-LEVLFQ↓G-6His-GST substrate. All these results were consistent with the flow cytometry results obtained from eYESS approach.

**Figure 2.**
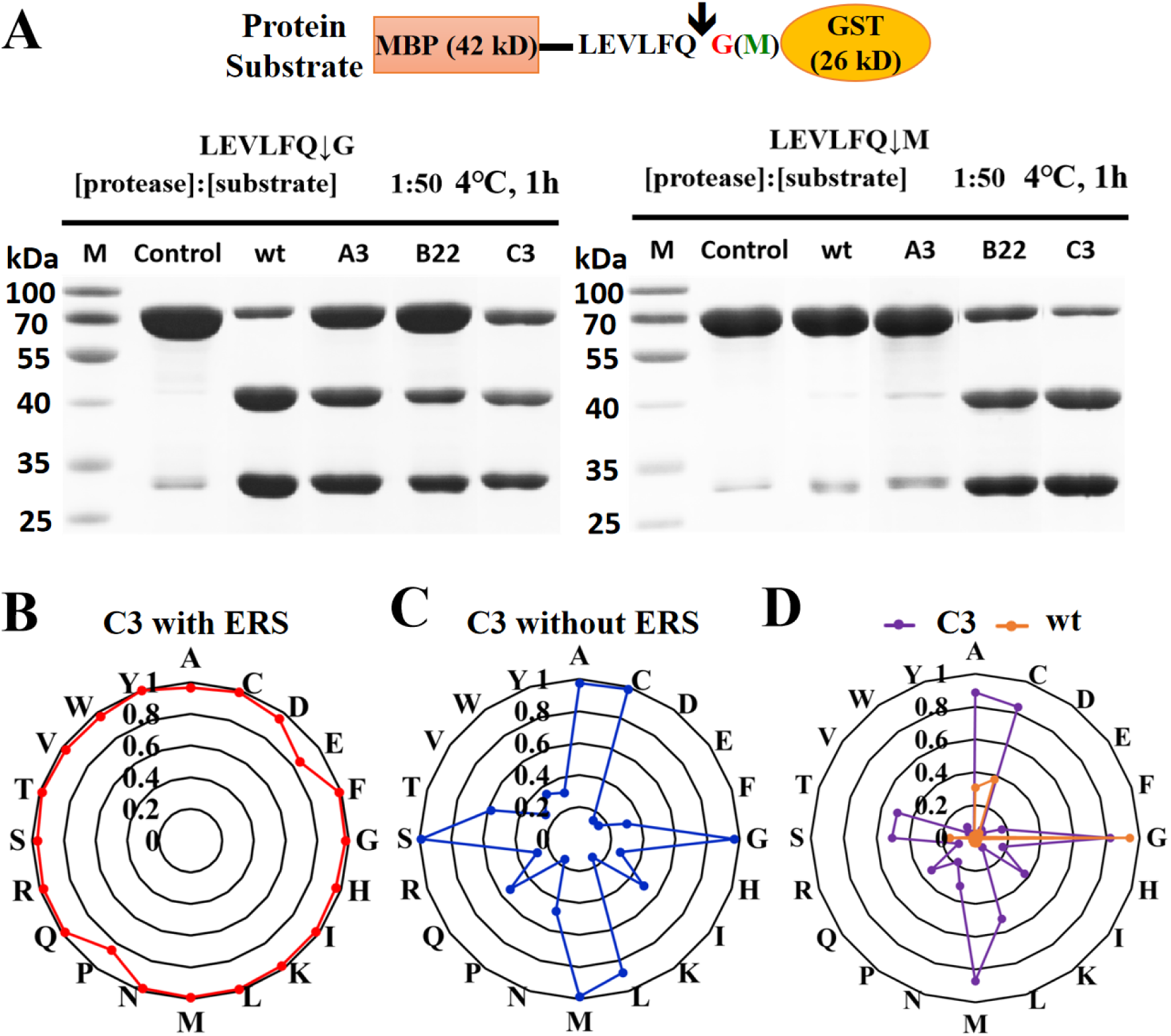
Characterization of the substrate specificity of HRV 3C-P variants at the P1’ site. A. Digestion of protein fusion substrates MBP-LEVLFQ↓G/M-6His-GST by purified HRV 3C-P and its variants, including A3, B22, and C3. The control only contains the substrate MBP-LEVLFQ↓G/M-6His-GST without protease. B. Substrate specificity of C3 at the P1’ site with ERS of FEHDEL at the C-terminus of both protease and substrate cassettes was determined by the YESS-PSSC approach. C. Substrate specificity of C3 at the P1’ site without ERS at the C-terminus of either protease or substrate cassette was determined by the YESS-PSSC approach. D. The *in vitro* protein substrate digestion assays of C3 and wt HRV 3C-P against substrates MBP-LEVLFQ↓X-6His-GST were carried out under the molar ratio of protease to substrate (1:10).

### 3.4. Characterization of the P1’ substrate specificity of the best variant C3

The P1’ substrate specificity of the C3 variant with the best performance was further characterized by evaluating its cleavage efficiency against all 20 different residues at P1’ site (LEVLFQ↓X, X can be any residue). Using the previously developed YESS-PSSC approach[20], the C3 variant presented effective cleavage against all 20 residues at P1’ site with efficiencies of 80%-99% under the condition of anchoring FEHDEL at the C-terminus of both protease and substrate cassettes (**Figure 2B, S2**). Comparably, under the same condition, only substrates with Gly, Cys, Ala, and Ser at P1’ site could be cleaved by wt HRV 3C protease with efficiencies of 98%, 76%, 71%, and 35%, respectively, while negligible cleavage were observed for other residues[20]. Under a more stringent condition that no ERS was anchored at the C-terminus of either protease or substrate cassette (**Figure 2C, S3**), C3 variant still showed high cleavage efficiency against most of the residues at P1’ site, except low cleavage efficiency of 10%-40% against residues with charged or aromatic side chains, including Asp, Glu, Phe, His, Lys, Pro, Arg, Val, Trp, and Tyr (**Figure 2C, S3**). In comparison, wt HRV 3C protease could only cleave Gly under this condition in YESS-PSSC[20].

These results were further confirmed through the *in vitro* protein substrate digestion assays against 20 different protein substrates (MBP-LEVLFQ↓X-6His-GST, X can be any residue) in a protease/substrate molar ratio of 1:10 at 4 [(**Figure 2D, S4**). C3 variant could recognize most of the residues at P1’ site, but presented lower activities against residues with charged or aromatic side chains (**Figure 2D, S4A**). While at the same condition, wt HRV 3C-P only showed cleavage efficiencies of 31%, 37%, 94%, and 16% against substrate having Ala, Cys, Gly, and Ser at the P1’ site, respectively (**Figure 2D, S4B**). It is worthy to point out that when we increase the protease/substrate molar ratio to 1:5 with a longer incubation time of 8 h at 4 [, C3 variant could effectively cleave protein substrates with charged or aromatic side chains including Phe, His, Lys, Arg, Val, Trp, and Tyr at P1’ site (**Figure S5**). Combined with the *in vitro* protein cleavage assay (**Figure 2D**) and in cell YESS-PSSC based cleavage efficiency evaluation (**Figure 2B** and **2C**), it could be concluded that the engineered C3 variant had a broaden substrate P1’ specificity, and could cleave substrates with different P1’ residues efficiently.

### 3.5. Kinetic analysis of the HRV 3C protease and C3 variant

To further compare the HRV 3C-P variants with wt HRV 3C-P, kinetic analysis against protein substrates were performed. The MBP-LEVLFQ↓X-6His-GFP (X can be any residue) substrates were used to mimic the protease cleavage against protein substrates. The cleaved substrates were analyzed by SDS-PAGE and the in-gel GFP fluorescence intensity was used to quantify the kinetic parameters of proteases (See Methods)[15]. It has to be pointed out that the *k_cat_*/*K_M_* values could only be well determined for C3 variant against 10 MBP-LEVLFQ↓X-6His-GFP substrates (X refers to Ala, Cys, Gly, Ile, Leu, Met, Asn, Gln, Ser, and Thr) (**Table 1, Figure S6**), because the cleavage against the other substrates were not effective enough to reach their maximum reaction rates. Due to the same reasons, the *k_cat_*/*K_M_* values for wt HRV 3C-P could only be well determined against substrates with Gly, Ala, Cys, and Ser at the P1’ site (**Table 1, Figure S7**). Our results showed that wt HRV 3C-P could efficiently cleave MBP-LEVLFQ↓G-6His-GFP substrate at 4 [with the *k_cat_*/*K_M_*value of 4.73 ± 0.01 mM^-1^·s^-1^ (**Table 1**). As to MBP-LEVLFQ↓X-6His-GFP (X refers to Ala, Cys, and Ser) substrates, the *k_cat_*/*K_M_*values decreased to 0.39-0.91 mM^-1^·s^-1^. Comparably, C3 variant showed slightly lower cleavage efficiency against MBP-LEVLFQ↓G-6His-GFP substrate at 4 [, with the *k_cat_*/*K_M_* value of 3.62 ± 0.12 mM^-1^·s^-1^. Nevertheless, it presented much higher cleavage efficiencies against other residues than those of wt HRV 3C protease, with most *k_cat_*/*K_M_*values ranging from 0.37 ± 0.01 to 3.72 ± 0.04 mM^-1^·s^-1^ (**Table 1**). It was also noticed that the C3 variant showed the highest *k_cat_*/*K_M_* value of 3.72 ± 0.04 mM^-1^·s^-1^ against MBP-LEVLFQ↓M-6His-GFP substrate, which might be because the libraries was sorted against the LEVLFQ↓M substrate. Although the kinetic parameters of C3 variant against MBP-LEVLFQ↓X-6His-GFP (X refers to Asp, Glu, Phe, His, Lys, Pro, Arg, Val, Trp, and Tyr) substrates could not be well determined, we still calculated the initial catalytic rate at the maximum concentration (360 μM) of these substrates. Compared to wt HRV 3C protease, C3 variant gave an approximately 10-fold higher initial catalytic rates against those substrates (**Table S3**).

**Table 1.**
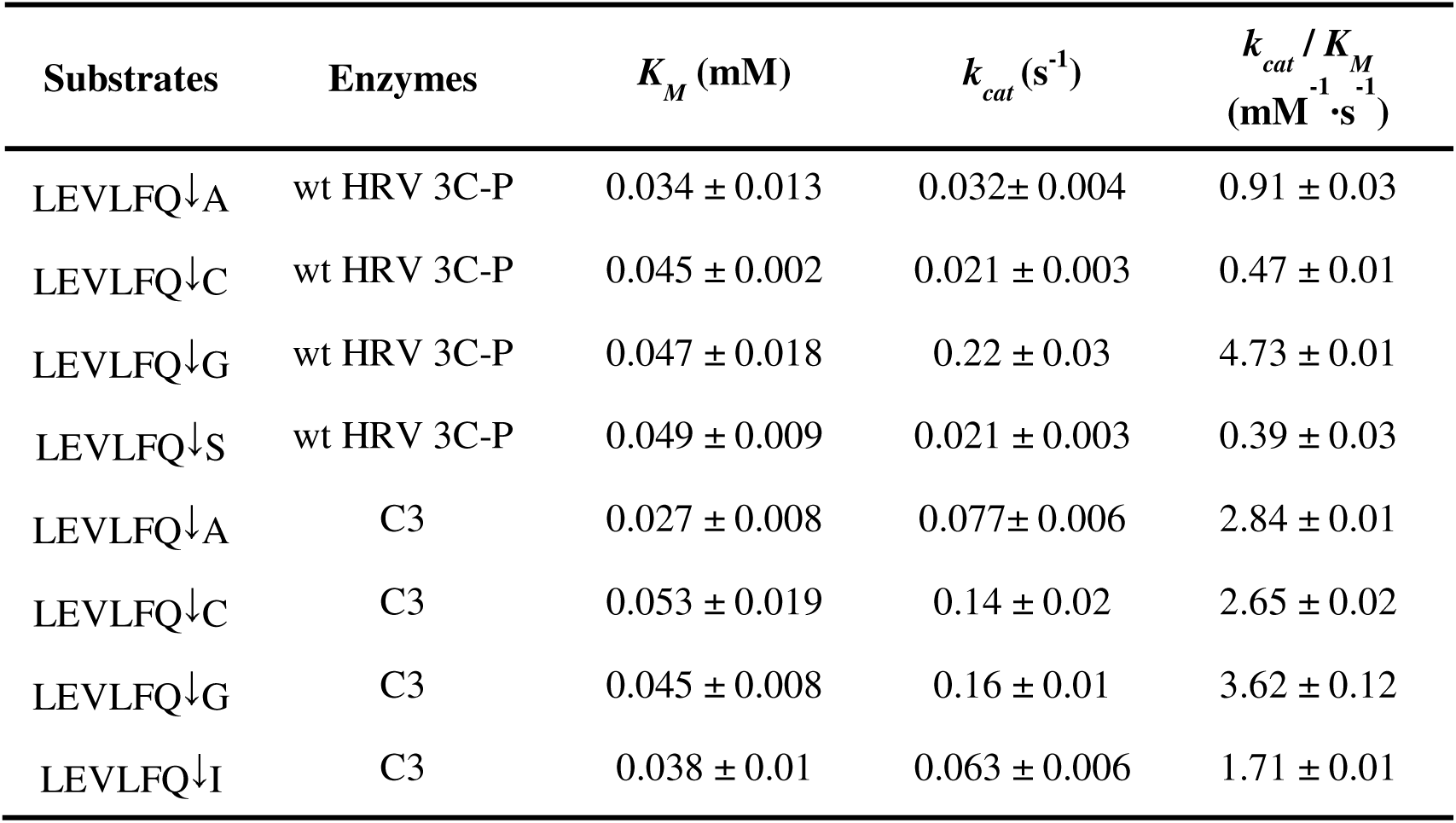
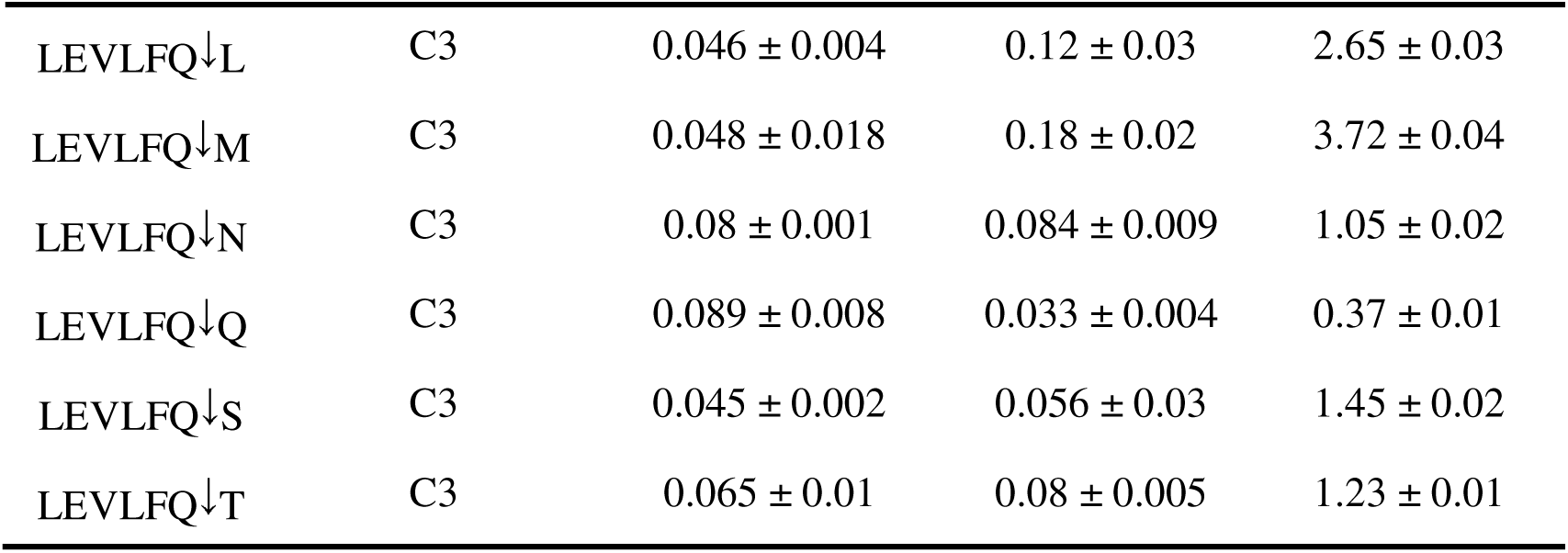
Kinetics data of wt HRV 3C-P and its variants.

| Substrates | Enzymes | $K_M$ (mM) | $k_{cat}$ ($\text{s}^{-1}$) | $k_{cat} / K_M$<br>($\text{mM}^{-1}\cdot\text{s}^{-1}$) |
| --- | --- | --- | --- | --- |
| LEVLFQ↓A | wt HRV 3C-P | $0.034 \pm 0.013$ | $0.032 \pm 0.004$ | $0.91 \pm 0.03$ |
| LEVLFQ↓C | wt HRV 3C-P | $0.045 \pm 0.002$ | $0.021 \pm 0.003$ | $0.47 \pm 0.01$ |
| LEVLFQ↓G | wt HRV 3C-P | $0.047 \pm 0.018$ | $0.22 \pm 0.03$ | $4.73 \pm 0.01$ |
| LEVLFQ↓S | wt HRV 3C-P | $0.049 \pm 0.009$ | $0.021 \pm 0.003$ | $0.39 \pm 0.03$ |
| LEVLFQ↓A | C3 | $0.027 \pm 0.008$ | $0.077 \pm 0.006$ | $2.84 \pm 0.01$ |
| LEVLFQ↓C | C3 | $0.053 \pm 0.019$ | $0.14 \pm 0.02$ | $2.65 \pm 0.02$ |
| LEVLFQ↓G | C3 | $0.045 \pm 0.008$ | $0.16 \pm 0.01$ | $3.62 \pm 0.12$ |
| LEVLFQ↓I | C3 | $0.038 \pm 0.01$ | $0.063 \pm 0.006$ | $1.71 \pm 0.01$ |
| LEVLFQ↓L | C3 | $0.046 \pm 0.004$ | $0.12 \pm 0.03$ | $2.65 \pm 0.03$ |
| LEVLFQ↓M | C3 | $0.048 \pm 0.018$ | $0.18 \pm 0.02$ | $3.72 \pm 0.04$ |
| LEVLFQ↓N | C3 | $0.08 \pm 0.001$ | $0.084 \pm 0.009$ | $1.05 \pm 0.02$ |
| LEVLFQ↓Q | C3 | $0.089 \pm 0.008$ | $0.033 \pm 0.004$ | $0.37 \pm 0.01$ |
| LEVLFQ↓S | C3 | $0.045 \pm 0.002$ | $0.056 \pm 0.03$ | $1.45 \pm 0.02$ |
| LEVLFQ↓T | C3 | $0.065 \pm 0.01$ | $0.08 \pm 0.005$ | $1.23 \pm 0.01$ |

### 3.6. Profiling the P1-P1’-P2’ substrate specificity of C3 variant using YESS-NGS approach

Previous studies indicated that HRV 3C-P has a high substrate specificity at P1 and P1’ sites, but can recognize most of residues at P2’ site[20]. Here, we profiled the substrate specificity of C3 variant at P1-P1’-P2’ positions using the combinatorial YESS-NGS approach[26]. A substrate library with saturation mutagenesis at the P1, P1’, and P2’ sites was generated and ligated into the pYSD-5 vector (**Table S1**) containing C3 variant. After 3 rounds of cell enrichment against high iFluor 647 and low FITC signals, the substrate DNA of the sorted library cells was extracted and further analyzed by next-generation sequencing (NGS) (**Figure 3**). Our results indicated that the C3 variant still kept very high substrate specificity against residues at P1 site in the LEVLFX↓XX, only recognizing Gln same as the wt HRV 3C-P. Not surprisingly, the NGS results confirmed that the C3 variant did not have strong residue preference at either P1’ or P2’ position, preferring to the Met and Pro at P1’ and P2’, respectively, with well accommodation of other residues. These results confirmed that engineered C3 only expanded its substrate specificity at P1’ position. Its merits, including high substrate stringency at P1 position and good residue tolerance at P2’ position, were sustained (**Figure 3**). In short, our original goal of expanding the substrate P1’ specificity of HRV 3C protease was achieved through engineering.

**Figure 3.**
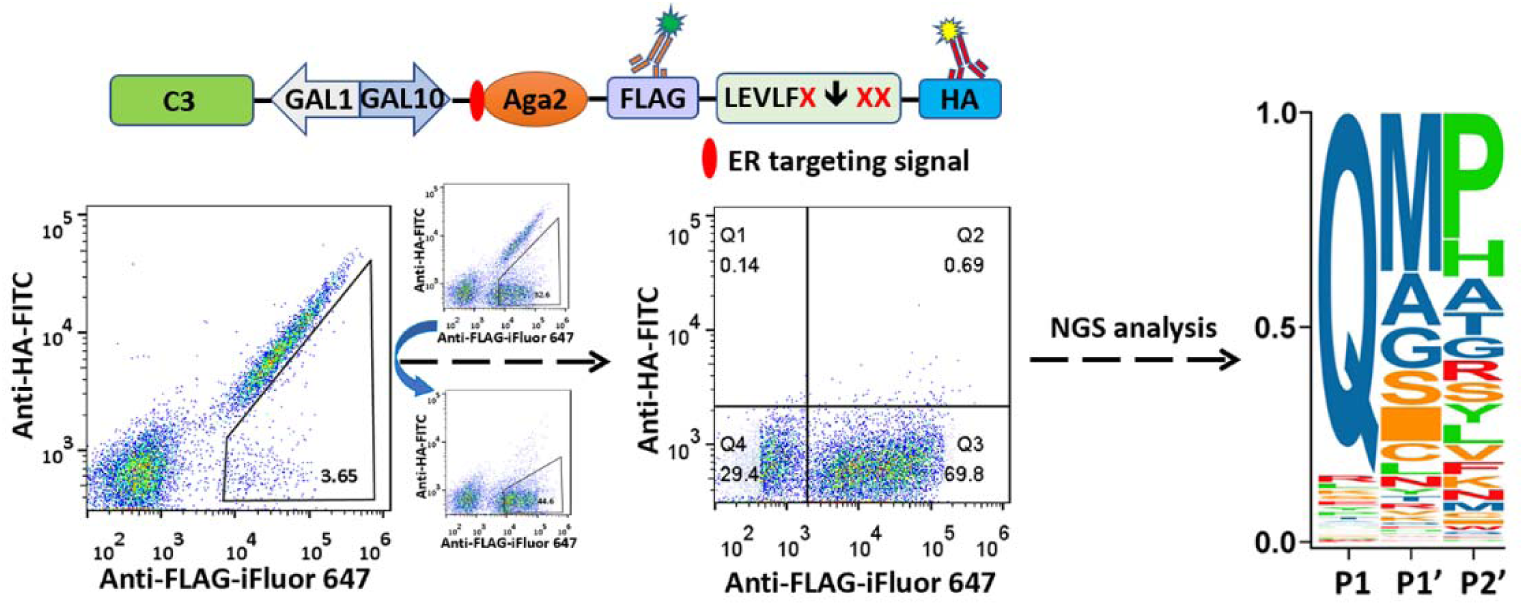
Characterization of the substrate P1-P1’-P2’ specificity of the C3 variant. Cell sorting and NGS analysis of C3 variant against substrate library LEVLFX↓XX. Three rounds of cell enrichment with high iFluor 647 fluorescence and low FITC fluorescence were carried out. For each round of sorted cells, plasmids were extracted for the DNA amplification, followed by NGS analysis. The substrate specificities of C3 at P1, P1’, and P2’ positions were shown in Seq2Logo results.

### 3.7. Structural characterization of the engineered variant C3

Structural simulation of HRV 3C-P and its variants were performed to understand the possible substrate recognition mechanisms. The structures of obtained HRV 3C-P variants were simulated by AlphaFold 2[29], followed by ZDOCK program in Discovery Studio (DS) with peptide substrate LEVLFQ↓M (**Figure 4**). As shown in **Figure 4A** and **Table S4**, the S1’ binding pocket in wt HRV 3C-P was very rigid and small, in which the distance between Thr143 and Lys22 at the entrance of the pocket was only 4.97Å, and the aromatic ring of the side chain at position F25 further squeezed the S1’ binding pocket. Thus only small residues such as Gly, Ala, Cys and Ser could be accommodated[20].

In comparison, we observed gradually changed S1’ binding pockets in the obtained HRV 3C-P variants. In A3 variant, the S1’ binding pocket became looser and exhibited a larger space due to the K22M, T143P, and Q145T mutations (**Figure 4B**). Comparing to the wt HRV 3C-P, the nearest atomic distances among residues 22, 106 and 143 in the S1’ pocket increased from 4.97 Å, 7.06 Å, and 6.75 Å (**Figure 4A, Table S4**) to 8.79 Å, 11.71 Å, and 10.60 Å, respectively (**Figure 4B, Table S4**). However, effective docking of Met into the S1’ binding pocket was still difficult, because the Phe25 showed a clear steric hindrance due to bulky side chain in the A3 variant. This issue was well solved in the B22 variant, in which Phe25 was mutated to Val with very small side chain to remove this steric hindrance. Besides, the B22 variant also caused a rearrangement of the S1’ binding pocket due to the mutations of Q145M, T143M, N106Y, K22L and F25V. Compared to A3 variant, the nearest atomic distances between residue 25 and residues 143, 106 and 22 increased from 9.92 Å, 7.10 Å, and 7.59 Å in A3 (**Figure 4C, Table S4**) to 11.99 Å, 10.29 Å, and 8.86 Å in B22, respectively (**Figure 4D, Table S4**). It could be speculated that this moderate change promotes the recognition efficiency of Met at the substrate P1’ site. Its expanded S1’ binding pocket conferred the ability of recognizing large side chain amino acids, such as Met. While on the other hand, the nearest atomic distances among residues 22, 106 and 143 decreased slightly from 8.79 Å, 11.71 Å, and 10.60 Å in A3 to 8.22 Å, 8.72 Å, and 9.63 Å in B22, respectively (**Table S4)**, which might help hold Met more tightly in B22.

**Figure 4.**
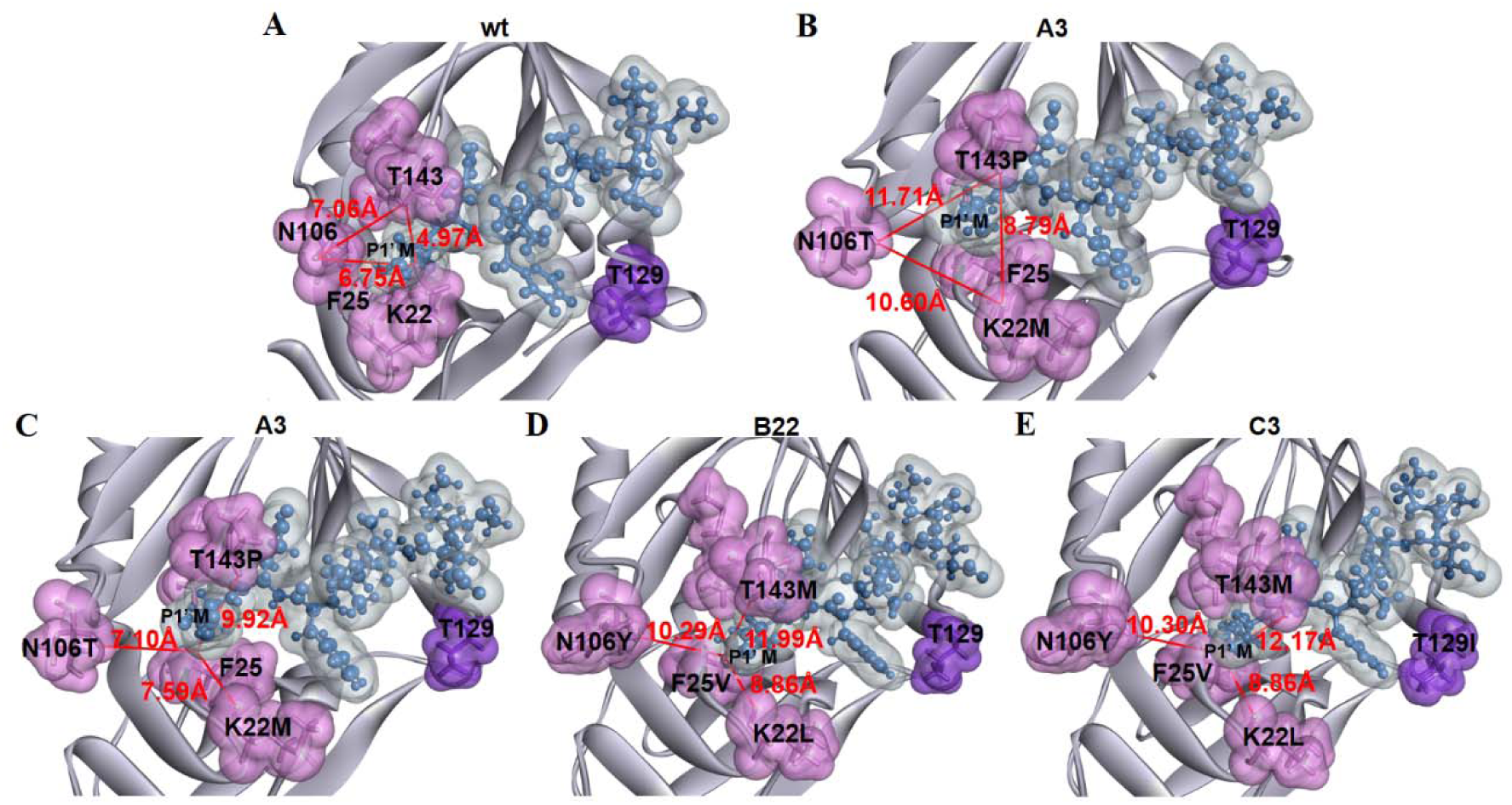
Molecular docking of HRV 3C-P and its variants with the substrate LEVLFQ**↓**M. (A-E) The LEVLFQ↓M substrate (blue) was docked in the wt HRV 3C-P and its variants (gray). The residues at 22, 25, 106,143, 144, and 145 position in the protease are labeled in pink spheres. The residue at 129 position is labeled in purple sphere. The nearest atomic distances among residues 22, 106 and 143 in wt HRV 3C-P (A) and A3 (B) are presented, respectively. Additionally, the nearest atomic distances between residues 25 and 143, 106 and 22 in A3 (C), B22 (D), and C3 (E) were also presented, respectively.

C3 variant was isolated from an error-prone library, which introduced an additional mutation, T129I, outside of the S1’ binding pocket. Based on the residue distance characterization, the size of S1’ binding pocket in B22 and C3 variants is very similar (**Figure 4E, Table S4**), indicating that the re-constituted S1’ binding pocket has been adjusted to an appropriate state to accommodate Met in P1’ position. It was speculated that the T129I mutation might help the correct orientation of substrate in the S1’ binding pocket. As shown in **Figure 5A**, Thr129 interacted with Leu126, Ser127, and Thr131 through strong H-bonds in wt HRV 3C-P. Similar H-bonds could be identified in A3 variant as well (**Figure 5B**). Comparably, the H-bonds between Thr129 and Thr131 in B22 variant (**Figure 5C**), and Ile129 and Ser127 in C3 variant (**Figure 5D**) are lost. These key H-bonds changes indicated that the mutation at 129 position might affect the interaction of its nearby residues with the substrate, thus affecting the substrate docking into the protease.

Detailed molecular docking analysis of wt HRV 3C-P and A3 indicated that two H-bonds formed between Asn125 and Ser127 with Val at the substrate P4 position to maintain its correct orientation in the S4 binding pocket (**Figure 5A and 5B**). However, the H-bond between the Asn125 and Val at the substrate P4 position is lost in B22 variant (**Figure 5C**). Very interestingly, this H-bond is recovered in C3 variant due to the T129I mutation, and a new H-bond is formed between the Ser127 and Phe at substrate P2 position (**Figure 5D**). We speculated that T129I mutation might help stabilize the Phe at P2 position and Val at P4 position in the S2 and S4 binding pocket of C3 variant, respectively. Therefore, C3 variant presented a nearly 1.3-fold improvement in cleavage efficiency in comparison with B22 variant (**Figure 2A**). It was also noticed that all the mutants showed a unmutated Gly at position 144 (**Table S2**), suggested its important role in maintaining the S1’ binding pocket by prompting flexibility of residues at 143 and 145 position.

**Figure 5.**
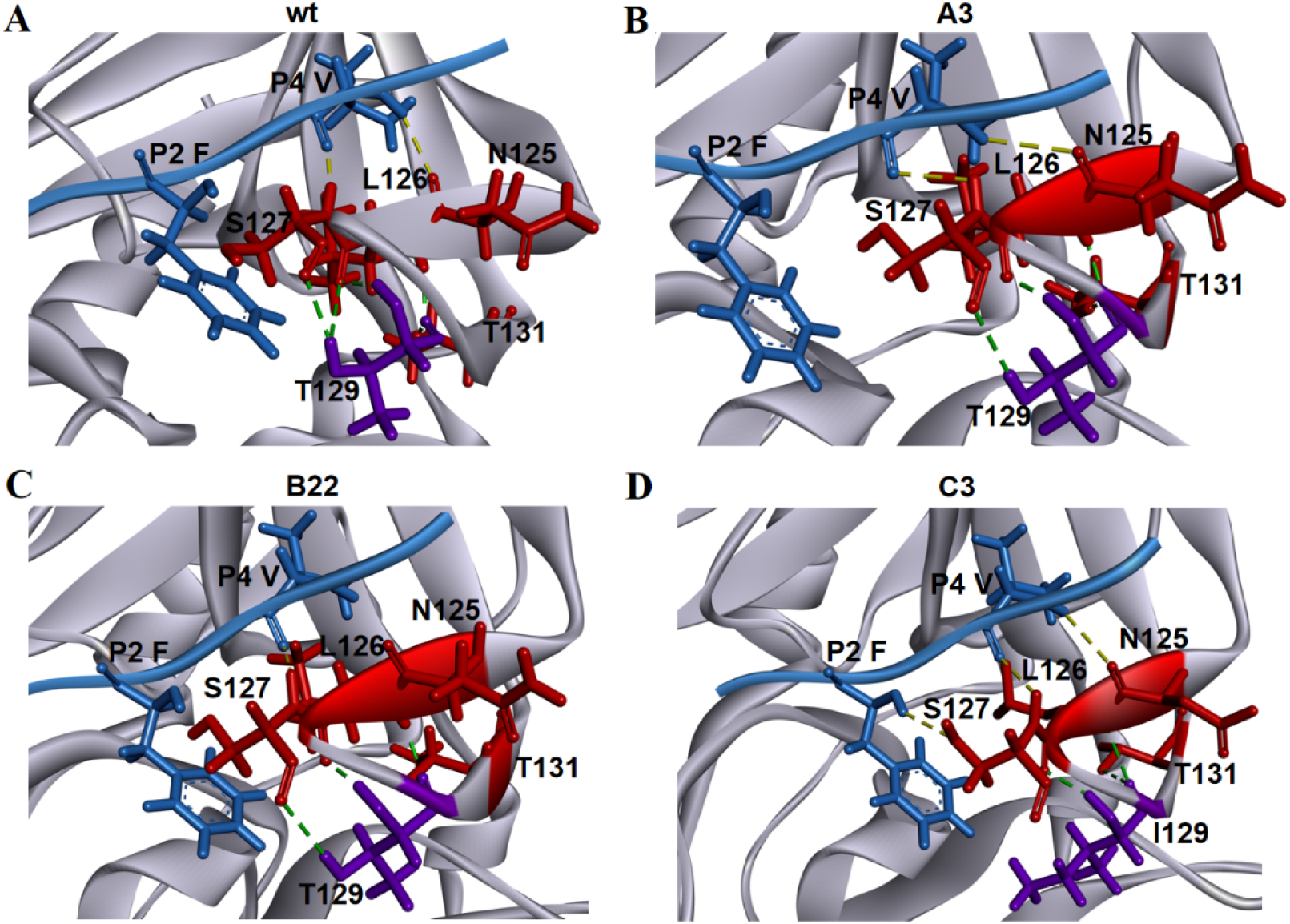
The possible interaction of HRV 3C-P and its variants with the substrate LEVLFQ**↓**M at the P2 and P4 position. The LEVLFQ↓M was docked in the wt HRV 3C-P (A) and its variants A3 (B), B22(C) and C3 (D). The substrate was shown in blue and proteases were shown in gray. The residues interacted with 129 residue (purple) were shown in red. The H-bonds between the 129 residue with Leu126, Ser127, and Thr131 in wt HRV 3C-P and its variants were shown in green. The H-bonds between the P2 and P4 residue of LEVLFQ↓M substrate with Asn125 and Ser127 in wt HRV 3C-P and its variants were shown in yellow.

## 4. Conclusion

In summary, an eYESS strategy was developed for HRV 3C protease engineering with altered P1’ substrate specificity. After analyzing a structured-guided library followed by an error-prone library, a C3 variant was obtained, showing attractive characteristics to keep high substrate specificity at P1 site with an expanded substrate specificity at P1’ site. Very different from wt HRV 3C protease, C3 variant presented a re-constructed S1’ pocket to recognize all 20 residues at the P1’ site, indicating its potentially broaden applications. Additionally, the simulated structure analysis of variants during the HRV 3C protease engineering also promotes our understanding of its substrate specificity mechanism. Moreover, since the YESS has been proved for other enzyme engineering[18], the eYESS strategy developed here would be generally useful for other proteases engineering.

## CRediT authorship contribution statement

Meng Mei, Xian Fan, Yu Zhou, Faying Zhang: performed the experiments. Meng Mei: analyzed the data and drafted the manuscript. Meng Mei, Guimin Zhang, and Li Yi: designed the experiments, wrote and edited the manuscript. All authors have read and agreed to the published version of the manuscript.

## Acknowledgments

This study was supported by the National Key Research and Development Program of China (No. 2018YFA0901100 to L. Yi and G. Zhang), the National Natural Science Foundation of China (No. 31870057 to L. Yi).

