## Supplemental file for "A Combinatorial Strategy for HRV 3C Protease Engineering to Achieve the N-terminal Free Cleavage"

### Supplementary Tables

#### Table S1. Constructs generated in this work

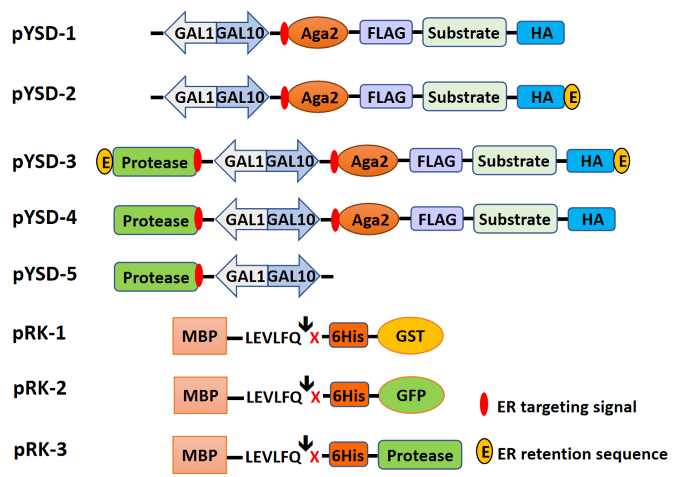

| **Construct** | | **ERS** | **Enzyme** | **Substrate** | **Construct** | | **ERS** | **Enzyme** | **Substrate** |
| --- | --- | --- | --- | --- | --- | --- | --- | --- | --- |
| pYSD-1 |  |  |  |  | pRK-1 |  |  |  |  |
|  | pYSD-1 | FEHDEL | / | LEVLFQ**↓**M |  | pRK-1-1 | / | / | LEVLFQ**↓**A |
| pYSD-2 |  |  |  |  |  | pRK-1-2 | / | / | LEVLFQ**↓**C |
|  | pYSD-2 | / | / | LEVLFQ**↓**M |  | pRK-1-3 | / | / | LEVLFQ**↓**D |
| pYSD-3 |  |  |  |  |  | pRK-1-4 | / | / | LEVLFQ**↓**E |
|  | pYSD-3-1 | FEHDEL | C3 | LEVLFQ**↓**A |  | pRK-1-5 | / | / | LEVLFQ**↓**F |
|  | pYSD-3-2 | FEHDEL | C3 | LEVLFQ**↓**C |  | pRK-1-6 | / | / | LEVLFQ**↓**G |
|  | pYSD-3-3 | FEHDEL | C3 | LEVLFQ**↓**D |  | pRK-1-7 | / | / | LEVLFQ**↓**H |
|  | pYSD-3-4 | FEHDEL | C3 | LEVLFQ**↓**E |  | pRK-1-8 | / | / | LEVLFQ**↓**I |
|  | pYSD-3-5 | FEHDEL | C3 | LEVLFQ**↓**F |  | pRK-1-9 | / | / | LEVLFQ**↓**K |
|  | pYSD-3-6 | FEHDEL | C3 | LEVLFQ**↓**G |  | pRK-1-10 | / | / | LEVLFQ**↓**L |
|  | pYSD-3-7 | FEHDEL | C3 | LEVLFQ**↓**H |  | pRK-1-11 | / | / | LEVLFQ**↓**M |
|  | pYSD-3-8 | FEHDEL | C3 | LEVLFQ**↓**I |  | pRK-1-12 | / | / | LEVLFQ**↓**N |
|  | pYSD-3-9 | FEHDEL | C3 | LEVLFQ**↓**K |  | pRK-1-13 | / | / | LEVLFQ**↓**P |
|  | pYSD-3-10 | FEHDEL | C3 | LEVLFQ**↓**L |  | pRK-1-14 | / | / | LEVLFQ**↓**Q |
|  | pYSD-3-11 | FEHDEL | C3 | LEVLFQ**↓**M |  | pRK-1-15 | / | / | LEVLFQ**↓**R |
|  | pYSD-3-12 | FEHDEL | C3 | LEVLFQ**↓**N |  | pRK-1-16 | / | / | LEVLFQ**↓**S |
|  | pYSD-3-13 | FEHDEL | C3 | LEVLFQ**↓**P |  | pRK-1-17 | / | / | LEVLFQ**↓**T |
|  | pYSD-3-14 | FEHDEL | C3 | LEVLFQ**↓**Q |  | pRK-1-18 | / | / | LEVLFQ**↓**V |
|  | pYSD-3-15 | FEHDEL | C3 | LEVLFQ**↓**R |  | pRK-1-19 | / | / | LEVLFQ**↓**W |
|  | pYSD-3-16 | FEHDEL | C3 | LEVLFQ**↓**S |  | pRK-1-20 | / | / | LEVLFQ**↓**Y |
|  | pYSD-3-17 | FEHDEL | C3 | LEVLFQ**↓**T | pRK-2 |  |  |  |  |
|  | pYSD-3-18 | FEHDEL | C3 | LEVLFQ**↓**V |  | pRK-2-1 | / | / | LEVLFQ**↓**A |
|  | pYSD-3-19 | FEHDEL | C3 | LEVLFQ**↓**W |  | pRK-2-2 | / | / | LEVLFQ**↓**C |
|  | pYSD-3-20 | FEHDEL | C3 | LEVLFQ**↓**Y |  | pRK-2-3 | / | / | LEVLFQ**↓**D |
| pYSD-4 |  |  |  |  |  | pRK-2-4 | / | / | LEVLFQ**↓**E |
|  | pYSD-4-1 | / | C3 | LEVLFQ**↓**A |  | pRK-2-5 | / | / | LEVLFQ**↓**F |
|  | pYSD-4-2 | / | C3 | LEVLFQ**↓**C |  | pRK-2-6 | / | / | LEVLFQ**↓**G |
|  | pYSD-4-3 | / | C3 | LEVLFQ**↓**D |  | pRK-2-7 | / | / | LEVLFQ**↓**H |
|  | pYSD-4-4 | / | C3 | LEVLFQ**↓**E |  | pRK-2-8 | / | / | LEVLFQ**↓**I |
|  | pYSD-4-5 | / | C3 | LEVLFQ**↓**F |  | pRK-2-9 | / | / | LEVLFQ**↓**K |
|  | pYSD-4-6 | / | C3 | LEVLFQ**↓**G |  | pRK-2-10 | / | / | LEVLFQ**↓**L |
|  | pYSD-4-7 | / | C3 | LEVLFQ**↓**H |  | pRK-2-11 | / | / | LEVLFQ**↓**M |
|  | pYSD-4-8 | / | C3 | LEVLFQ**↓**I |  | pRK-2-12 | / | / | LEVLFQ**↓**N |
|  | pYSD-4-9 | / | C3 | LEVLFQ**↓**K |  | pRK-2-13 | / | / | LEVLFQ**↓**P |
|  | pYSD-4-10 | / | C3 | LEVLFQ**↓**L |  | pRK-2-14 | / | / | LEVLFQ**↓**Q |
|  | pYSD-4-11 | / | C3 | LEVLFQ**↓**M |  | pRK-2-15 | / | / | LEVLFQ**↓**R |
|  | pYSD-4-12 | / | C3 | LEVLFQ**↓**N |  | pRK-2-16 | / | / | LEVLFQ**↓**S |
|  | pYSD-4-13 | / | C3 | LEVLFQ**↓**P |  | pRK-2-17 | / | / | LEVLFQ**↓**T |
|  | pYSD-4-14 | / | C3 | LEVLFQ**↓**Q |  | pRK-2-18 | / | / | LEVLFQ**↓**V |
|  | pYSD-4-15 | / | C3 | LEVLFQ**↓**R |  | pRK-2-19 | / | / | LEVLFQ**↓**W |
|  | pYSD-4-16 | / | C3 | LEVLFQ**↓**S |  | pRK-2-20 | / | / | LEVLFQ**↓**Y |
|  | pYSD-4-17 | / | C3 | LEVLFQ**↓**T | pRK-3 |  |  |  |  |
|  | pYSD-4-18 | / | C3 | LEVLFQ**↓**V |  | pRK-3-1 | / | wt HRV 3C-P | LEVLFQ**↓**G |
|  | pYSD-4-19 | / | C3 | LEVLFQ**↓**W |  | pRK-3-2 | / | A3 | LEVLFQ**↓**M |
|  | pYSD-4-20 | / | C3 | LEVLFQ**↓**Y |  | pRK-3-3 | / | B22 | LEVLFQ**↓**M |
| pYSD-5 |  | / | C3 | / |  | pRK-3-4 | / | C3 | LEVLFQ**↓**M |

#### Table S2. HRV 3C-P variants obtained from eYESS

| **Variants** | **K22** | **F25** | **N106** | **T129** | **A140** | **T143** | **G144** | **Q145** |
| --- | --- | --- | --- | --- | --- | --- | --- | --- |
| A1 | R | F | E | T | A | F | G | M |
| A2 | P | F | N | T | A | T | G | Q |
| A3 | M | F | T | T | A | P | G | T |
| A4 | V | F | G | T | A | F | G | Q |
| A5 | W | F | G | T | A | T | G | Q |
| A6 | Q | F | G | T | A | T | G | Q |
| A7 | W | F | N | T | A | T | G | Q |
| A8 | V | F | G | T | A | F | G | F |
| A9 | K | F | G | T | A | W | G | W |
| A10 | L | F | V | T | A | F | G | M |
| A11 | W | F | G | T | A | P | G | D |
| A12 | M | F | N | T | A | T | G | Q |
| A13 | Q | F | G | T | A | Y | G | Q |
| A14 | R | F | R | T | A | W | G | Y |
| A25 | C | F | N | T | A | T | G | Q |
| A26 | P | F | R | T | A | F | G | H |
| A31 | G | F | C | T | A | P | C | F |
| A32 | L | F | N | T | A | T | G | Q |
| A35 | R | F | G | T | A | F | G | F |
| A43 | P | F | G | T | A | M | G | Q |
| B5 | L | V | K | T | A | P | G | T |
| B7 | R | V | N | T | A | T | G | Q |
| B12 | Q | C | C | T | A | P | G | T |
| B16 | R | L | G | T | A | P | G | T |
| B20 | L | V | N | T | A | T | G | Q |
| B21 | R | V | K | T | A | P | G | M |
| B22 | L | V | Y | T | A | M | G | M |
| B23 | A | F | F | T | A | A | G | M |
| B24 | L | V | W | T | A | P | G | C |
| B27 | L | V | N | T | A | F | G | M |
| C3 | L | V | Y | I | A | M | G | M |
| C9 | L | V | Y | T | S | M | G | M |

#### Table S3. The initial catalytic rates of wt HRV 3C-P and C3 variant against the protein substrates at their maximum protein concentration of 360 μM

| **Substrate** | **Enzyme** | ***k*_,_ s^-1^** |
| --- | --- | --- |
| LEVLFQ↓D | wt HRV 3C-P | 3.1 ± 0.2 × 10^-4^ |
| LEVLFQ↓E | wt HRV 3C-P | 6.9 ± 0.3 × 10^-4^ |
| LEVLFQ↓F | wt HRV 3C-P | 3.6 ± 0.4 × 10^-4^ |
| LEVLFQ↓H | wt HRV 3C-P | 3.3 ± 0.3 × 10^-4^ |
| LEVLFQ↓I | wt HRV 3C-P | 4.2 ± 0.4 × 10^-4^ |
| LEVLFQ↓K | wt HRV 3C-P | 1.2 ± 0.2 × 10^-4^ |
| LEVLFQ↓L | wt HRV 3C-P | 6.2 ± 0.1 × 10^-4^ |
| LEVLFQ↓M | wt HRV 3C-P | 2.1 ± 0.3 × 10^-4^ |
| LEVLFQ↓N | wt HRV 3C-P | 2.5 ± 0.1 × 10^-4^ |
| LEVLFQ↓P | wt HRV 3C-P | 2.8 ± 0.2 × 10^-4^ |
| LEVLFQ↓Q | wt HRV 3C-P | 5.1 ± 0.2 × 10^-4^ |
| LEVLFQ↓R | wt HRV 3C-P | 1.1 ± 0.2 × 10^-4^ |
| LEVLFQ↓T | wt HRV 3C-P | 1.1 ± 0.1 × 10^-4^ |
| LEVLFQ↓V | wt HRV 3C-P | 1.6 ± 0.2 × 10^-4^ |
| LEVLFQ↓W | wt HRV 3C-P | 5.1 ± 0.6 × 10^-4^ |
| LEVLFQ↓Y | wt HRV 3C-P | 0.8 ± 0.1 × 10^-4^ |
| LEVLFQ↓D | C3 | 5.4 ± 0.1 × 10^-3^ |
| LEVLFQ↓E | C3 | 1.1 ± 0.2 × 10^-3^ |
| LEVLFQ↓F | C3 | 9.1 ± 0.1 × 10^-3^ |
| LEVLFQ↓H | C3 | 3.7 ± 0.2 × 10^-3^ |
| LEVLFQ↓K | C3 | 2.7 ± 0.2 × 10^-3^ |
| LEVLFQ↓P | C3 | 3.1 ± 0.5 × 10^-3^ |
| LEVLFQ↓R | C3 | 1.3 ± 0.1 × 10^-3^ |
| LEVLFQ↓V | C3 | 2.6 ± 0.3 × 10^-4^ |
| LEVLFQ↓W | C3 | 1.9 ± 0.2 × 10^-3^ |
| LEVLFQ↓Y | C3 | 1.6 ± 0.3 × 10^-3^ |

#### Table S4. The size of S1’ binding pocket in wt HRV 3C-P and its variants

| wt | Distance (Å) | A3 | Distance (Å) | B22 | Distance (Å) | C3 | Distance (Å) |
| --- | --- | --- | --- | --- | --- | --- | --- |
| T143-K22 | 4.97 | P143-M22 | 8.79 | M143-L22 | 8.22 | M143-L22 | 8.35 |
| T143-N106 | 7.06 | P143-T106 | 11.71 | M143-Y106 | 8.72 | M143-Y106 | 8.82 |
| N106-K22 | 6.75 | T106-M22 | 10.60 | Y106-L22 | 9.63 | Y106-L22 | 9.67 |
| T143-F25 | 9.89 | P143-F25 | 9.92 | M143-V25 | 11.99 | M143-V25 | 12.17 |
| N106-F25 | 9.74 | T106-F25 | 7.10 | Y106-V25 | 10.29 | Y106-V25 | 10.30 |
| K22-F25 | 6.17 | M22-F25 | 7.59 | L22-V25 | 8.86 | L22-V25 | 8.86 |

### Supplementary Figures

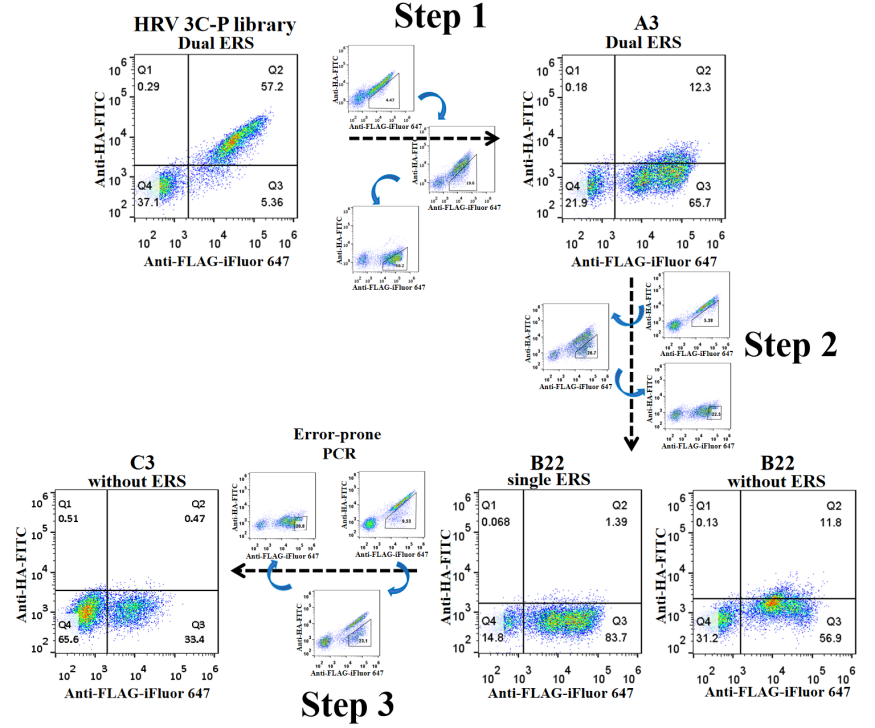

#### Figure S1. Cell sorting of HRV 3C-P libraries

In the first step, dual ERSs of FEHDEL were used at both of the protease library and its substrate. In the second step, the ERS of FEHDEL at the C terminus of the substrate was removed. For the third step, the ERS of FEHDEL at the C terminus of the protease library was also removed. Three rounds of cell enrichment with high iFluor 647 fluorescence and low FITC fluorescence were carried out throughout the total cell screening process.

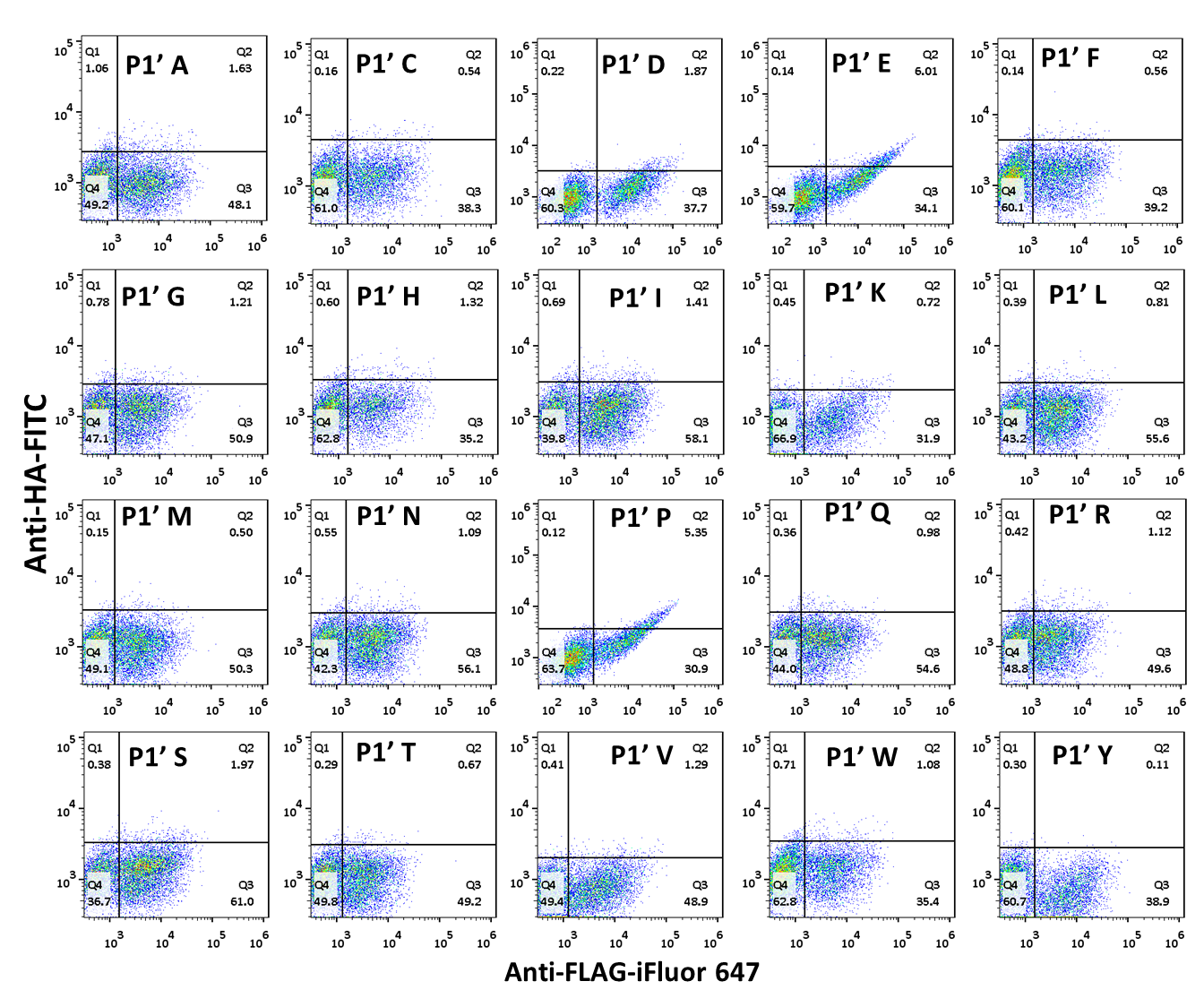

#### Figure S2. Quantitative analysis of the substrate specificity of C3 variant at P1’ position with double ERS of FEHDEL

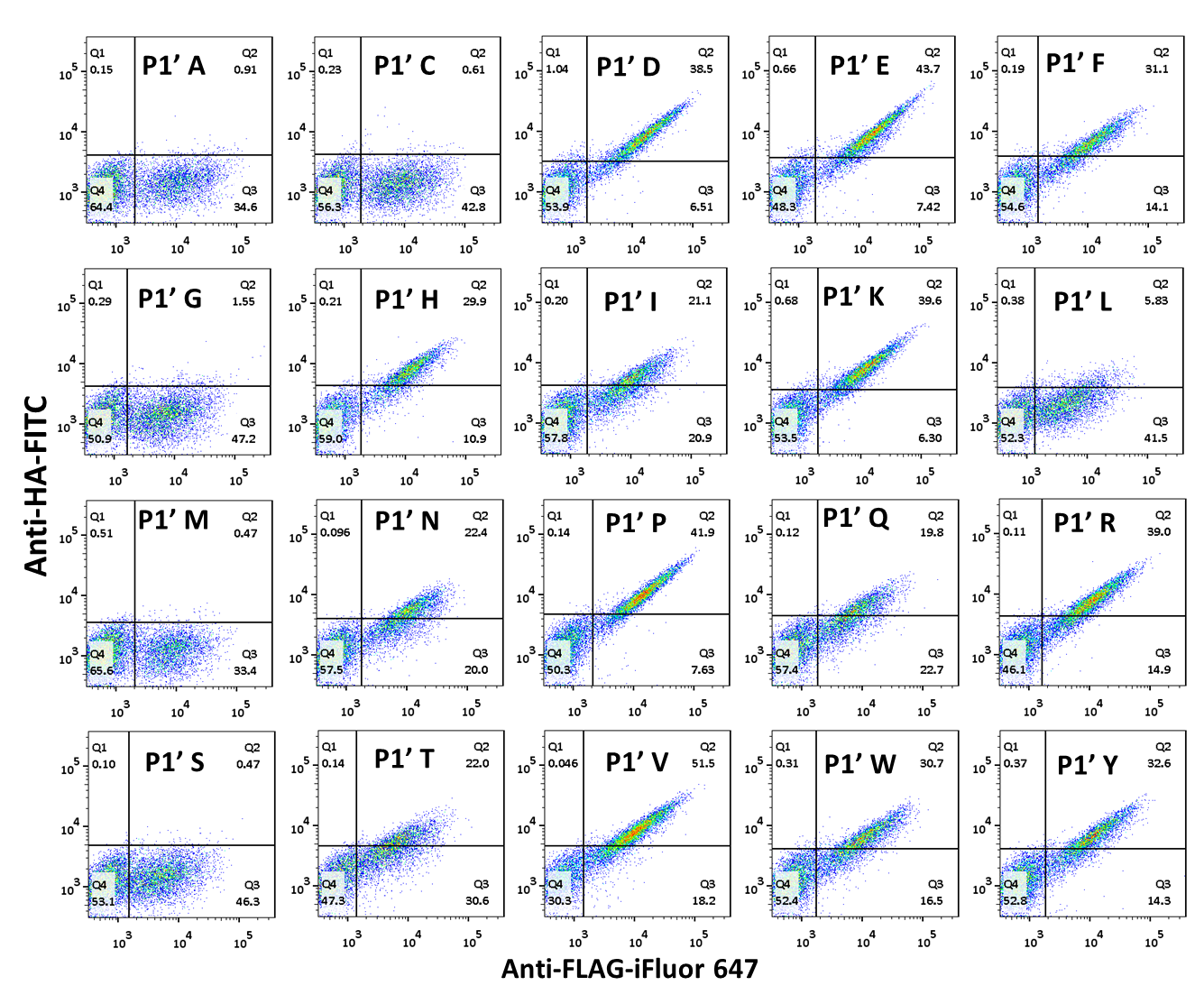

#### Figure S3. Quantitative analysis of the substrate specificity of C3 variant at P1’ position without ERS

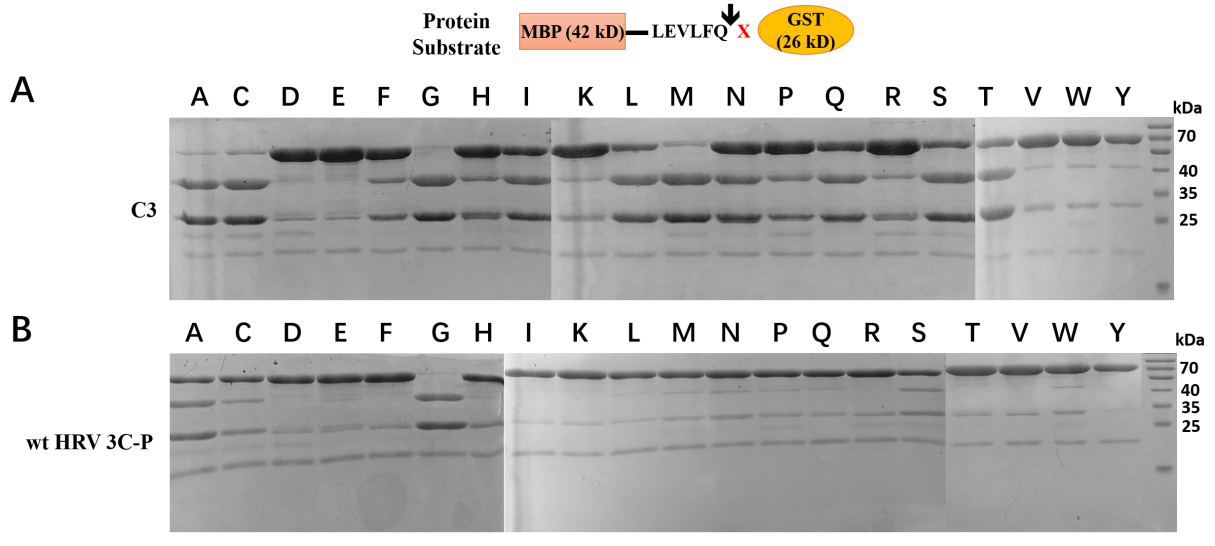

#### Figure S4. In vitro protein substrate digestion assays of wt HRV 3C-P and C3 variant against 20 protein substrates

The reaction was performed at 4 ℃ for 1 h with a ratio of 1:10 for protease to substrate.

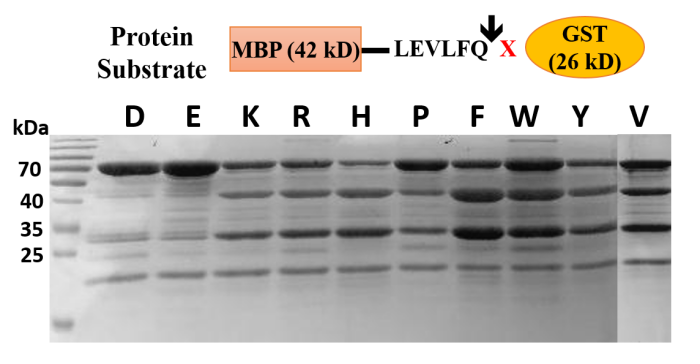

#### Figure S5. In vitro protein substrate digestion assay of C3 against substrates MBP-LEVLFQ↓D/E/K/R/H/P/F/W/Y/V-6His-GST

The reaction was performed at 4 ℃ for 8 h with a ratio of 1:5 for protease to substrate.

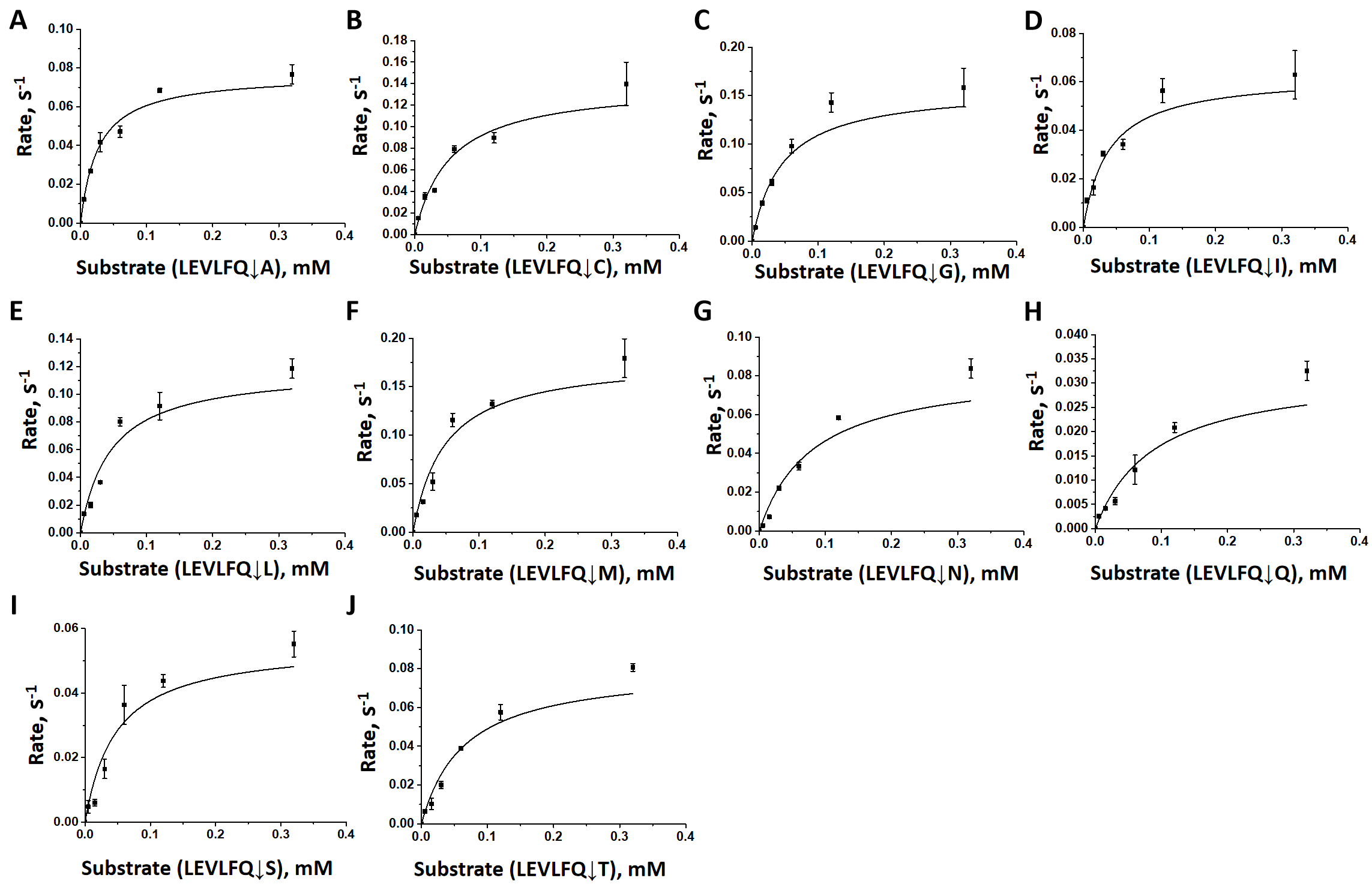

#### Figure S6. Michaelis-Menten plots for C3 variant against different protein substrates

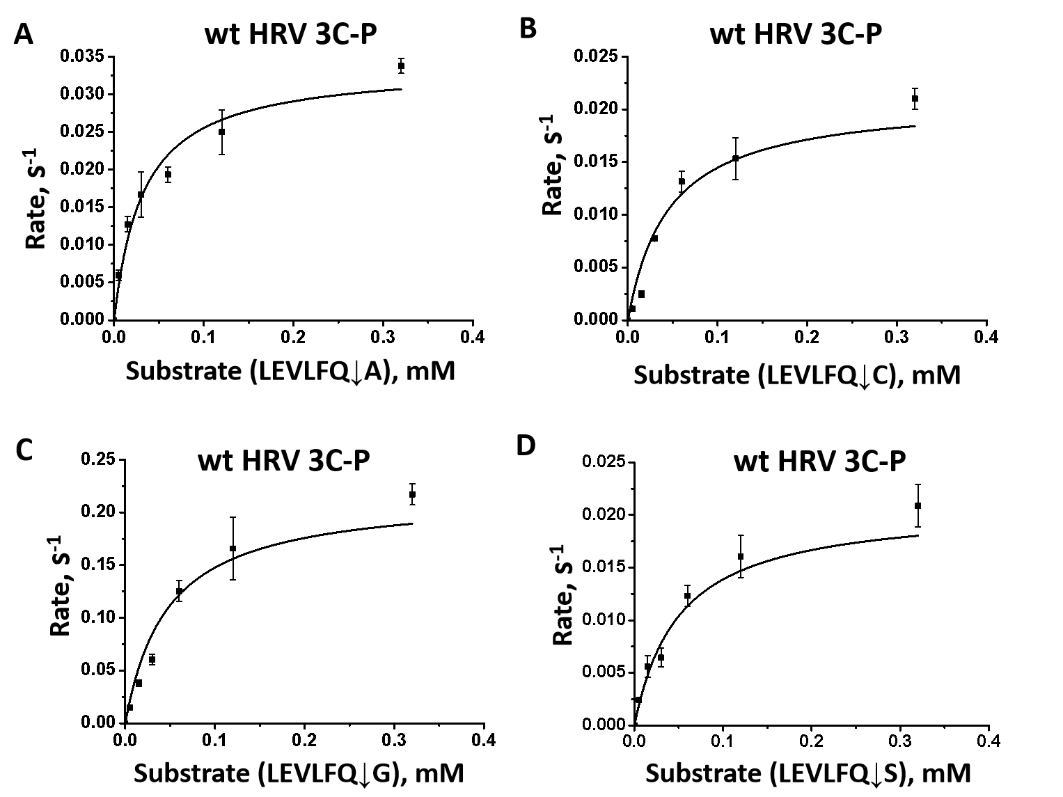

#### Figure S7. Michaelis-Menten plots for wt HRV 3C-P against different protein substrates
